# Genome-wide analysis of drug resistant Mycobacterium tuberculosis isolates causing pulmonary and extrapulmonary tuberculosis in Russia

**DOI:** 10.1101/486746

**Authors:** Ekaterina Chernyaeva, Mikhail Rotkevich, Ksenia Krasheninnikova, Alla Lapidus, Dmitriy E. Polev, Natalia Solovieva, Viacheslav Zhuravlev, Piotr Yablonsky, Stephen J. O’Brien

## Abstract

*Mycobacterium tuberculosis* is a highly studied pathogen due to public health importance. Despite progress in *M.tuberculosis* genome diversity analysis there remain insufficient data on genome analysis of *M.tuberculosis* strains associated with pulmonary vs. extrapulmonary TB (PTB or XPTB respectively) tissue localization. Here we conduct comparative analysis of whole-genome sequence (WGS) for clinical *M.tuberculosis* strains collected from patients with PTB (n=72) and XPTB (n=73) localization. We further analyze the incidence of point mutations widely used for drug resistance detection in laboratory practice.

*M.tuberculosis* isolates were collected from patients with varying clinical status from 2007 to 2014 in Russia. Bacterial DNA was extracted and sequenced using MiSeq platform (Illumina).

WGS data allowed identifying *M.tuberculosis* substrains associated with distinctions in the occurrence in PTB v s. XPTB cases. There occurred little statistically significant specific DNA variants or genotypes diagnostic of tissue distribution, however phylogenetic analyses did reveal *M.tuberculosis* genetic substrains associated with TB localization. XPTB was associated with Beijing CAO, A and 4.8 groups, while PTB localization was associated with group LAM (4.3). Further, XPTB strain in some cases showed elevated drug resistance patterns relative to PTB isolates. HIV is significantly associated with the development of XPTB in the Beijing B0/W148 group and among unclustered Beijing isolates.

This research analysis pinpointed genomic markers identified in XPTB and PTB with detailed characterization of drug-resistance markers of Russian *M.tuberculosis* isolates. We suggest that further comprehensive analysis of bacterial and human biological signatures might allow for better understanding consistent pattern of XPTB development.

**Summary:** This paper gives genome-wide characteristics of *Mycobacterium tuberculosis* strains collected from pulmonary and extrapulmonary tuberculosis patients:

- *M.tuberculosis* genetic strains associated with TB tissue localization,

- influence of HIV co-infection on TB dissemination;

- mutations associated with *M.tuberculosis* drug resistance.

## Introduction

*Mycobacterium tuberculosis* is one of the most widespread and studied pathogens across the globe. According to WHO estimation Russia had about 94,000 new TB cases (case rate 66 per 100 000) and 13700 TB deaths in 2016 with 27% of new TB cases displaying multiple drug resistance or resistance to rifampicin (MDR/RR-TB) (1). The main clinical form of tuberculosis (TB), pulmonary tuberculosis (PTB) is considered to be the most epidemically dangerous localization of the disease. For many years the proportion of patients with XPTB has remained constant with variation < 3% in Russia (2). Relatively low rate of XPTB in Russia could be explained by fact that TB of intrathoracic lymph nodes and tuberculous pleurisy are generally not counted in XPTB statistics. Delayed detection of XPTB leads to a high proportion of chronic TB forms and to a high level of disability among patients. The most common localization of XPTB is an osteoarticular localization, which is detected in 35.7% cases of all XPTB cases (2). TB localization differential may be influenced by differences of the host the immune status and genome features, as well as with the biological or genetic differences of the pathogen. In this study we compare the genomic landscape of clinical *M.tuberculosis* strains collected from patients with PTB and XPTB localization based on whole-genome sequencing data. XPTB *M.tuberculosis* WGS data were previously described by our team (3).

## Methods

A total of 72 PTB and 73 XPTB isolates were collected between 2007 to 2014 from 40 different regions of the Russia (fig. 1, supplementary table 1). Isolates were selected randomly from the *M. tuberculosis* strains collection of St. Petersburg Research Institute of Phthisiopulmonology. Eighteen isolates received from XPTB localization were collected from patient with generalized TB (disseminated form of TB). All TB patients in Russia are tested for HIV. The majority of studies isolates (n=120) were obtained from HIV-negative patients and 25 – from HIV-infected patients.

**Fig 1.**
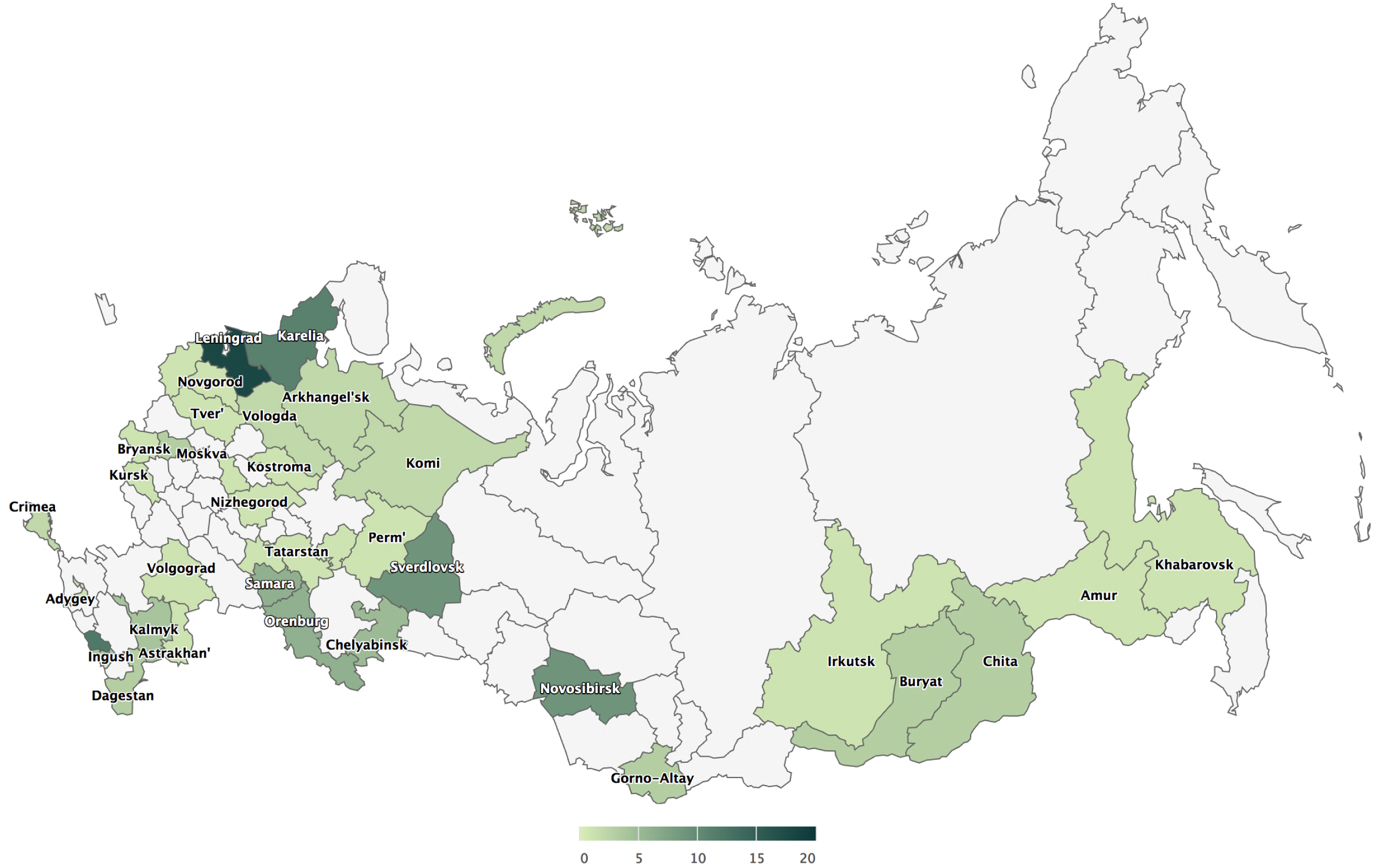
Geographical regions of Russia where *M. tuberculosis* patients were originated are highlighted with green color. The number of isolates collected in each region corresponds to the level of color saturation.

Bacterial isolates were cultured from pulmonary and extrapulmonary clinical material. The susceptibility of *M.tuberculosis* cultures to isoniazid (INH), rifampicin (RIF), streptomycin (SM), ethambutol (EMB), pyrazinamide (PZA), ethionamide (ETH), ofloxacin (OFL), kanamycin (KM), amikacin (AM), cycloserine (CS), capreomycin (CM) and para-aminosalicylic acid (PAS) was detected using WHO recommendations (4).

*M.tuberculosis* DNA was extracted using lysis with proteinase K and CTAB with further phenol/chloroform extraction and alcohol precipitation (5). Bacterial DNA was subjected to whole-genome sequencing (WGS) using MiSeq platform (Illumina). *M.tuberculosis* WGS data were deposited in the NCBI SRA (PRJNA352769).

The quality of M.tuberculosis sequence reads was evaluated using FastQC (http://www.bioinformatics.babraham.ac.uk/projects/fastqc/). Sequence reads were processed using Trimmomatic (6). We aligned sequenced reads to the reference genome and called variants (single-nucleotide polymorphisms [SNPs] and short insertions/deletions) by using bioinformatics software: bowtie2 (http://bowtie-bio.sourceforge.net/bowtie2/index.shtml); SAMtools (http://samtools.sourceforge.net); VCFtools (http://vcftools.sourceforge.net); and FreeBayes (https://github.com/ekg/freebayes). We used mutations that had q-scores ≥20 for comprehensive analysis. We used concatenated SNPs for phylogenetic analysis by using the GTRCAT (general time-reversible model with rate heterogeneity accommodated by using discrete rate categories) maximum-likelihood algorithm from the RAxML software package (7) to calculate an approximation model and 100 bootstrap replications. To avoid misalignments, we annotated SNPs in repetitive genome regions and in genes encoding proteins that contain proline-glutamate or proline-proline-glutamate motifs and filtered them from analysis. We used PhyTB (8) and SpoTyping tools (9) for phylogenetic classification of M. tuberculosis genomes. R commander package for R (https://socialsciences.mcmaster.ca/jfox/Misc/Rcmdr/) and Python scripts were used for statistical analysis using Fisher exact tests (FET) for significant statistical association.

## Results

### *M.tuberculosis* SNVs, genotypes and strains association with TB localization

Phylogenetic analysis discriminated two large lineages among sequenced *M.tuberculosis* isolates – 2 and 4 (fig. 2). Lineage 4 was represented by four monophyletic clusters which belonged to phylogenetic groups 4.1, 4.2, 4.3 and 4.8 based on PhyTB classification (9) and one isolate belonged to 4.4 phylogenetic groups. Major lineage 2 (Beijing) was represented by isolates belonged to 2.2.1 group based on PhyTB classification. Phylogenetic analysis based on 8673 SNVs allowed distinguishing several sub-clusters in lineage 2. Analysis of DNA markers such as regions of difference RD105, RD207, RD181 and mutT2 and mutT4 genes (10) allowed to classify “ancient” and “modern” sub-lineages within Beijing genotype (fig. 2). The largest sub-group within Beijing clade was a group of strains that belonged to B0/W148 genetic lineage (fig. 2). B0/W148 genetic cluster is characterized by a specific IS*6110* insertion in genome position 2,982,598 in the *Rv2664*-*Rv2665* intergenic region (11). Besides Beijing B0/W148 sublineage we identified clusters belonging to Beijing sublineages Central Asia (Beijing CAO) and Central Asia Clade A (Beijing A) based on SNV classification published by Shitikov et al. (12).

**Fig 2.**
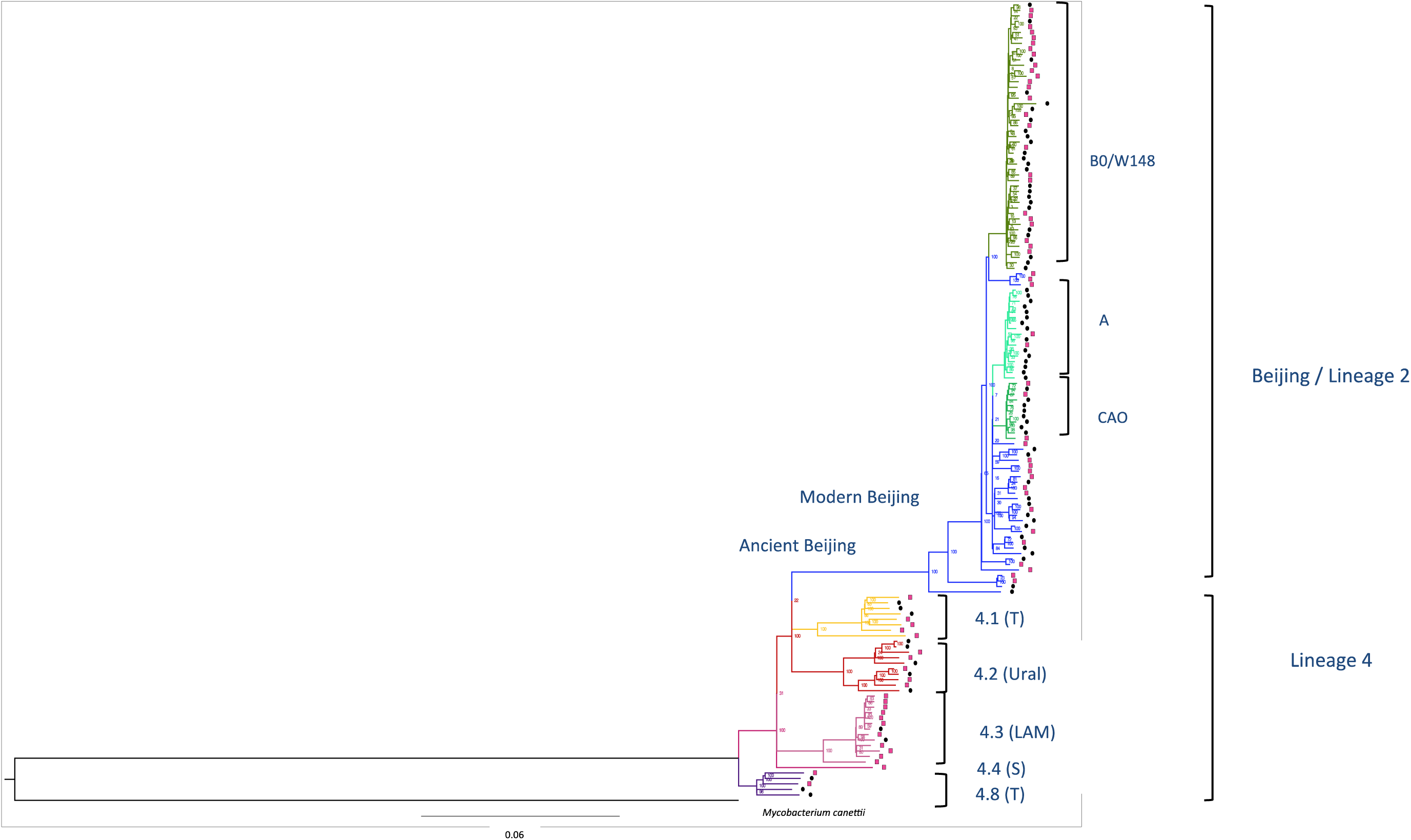
Phylogenetic analysis of *M. tuberculosis* isolates obtained from patients with PTB and XPTB. Phylogenetic groups were identified using maximum-likelihood approach. Isolates obtained from patients with PTB are marked using pink square symbol and from XPTB – with black square symbol. Beijing genetic group (lineage 2), colored with the blue, could be discriminated on 3 subclusters (B0/W148, Clades A and CAO marked with different shades of green) and a group of unclustered genotypes. Lineage 4 is represented by Ural (red), LAM (pink color) and T genetic families, one isolate belongs to genetic group 4.4, spoligotype family S. Detailed information on *M. tuberculosis* genetic lineages, microbiological and clinical data are available in Supplementary table 1.

The Beijing genetic group was predominant among both pulmonary and extrapulmonary isolates (Table 1), however, Beijing isolates were significantly more frequent among XPTB patients (82.19%, n=60) than among PTB patients (66.67%, n=48, p=0.03721). A large proportion of *M.tuberculosis* Beijing isolates (45.37%, n=59) belonged to B0/W148 sub-cluster. The Beijing B0/W148 group was equally represented in PTB vs. XPTB groups of bacterial isolates, while the frequency of Beijing sublineages CAO and Clade A was much higher among XPTB isolates. Beijing CAO strains was identified among 4.19% (n=3) PTB and 10.96% (n=8) XPTB cases; Beijing Clade A was detected among 4.19% (n=3) PTB and 19.18% (n=14) XPTB cases. Comparative analysis of XPTB cases caused by bacterial isolates from 4.8 and 4.3 genetic groups revealed significant differences in these clusters: *M.tuberculosis* isolates from 4.8 subcluster caused XPTB more often than isolates from 4.3 subcluster (p=0.02171). *M.tuberculosis* isolates from phylogenetic subcluster 4.3 was more often detected among PTB cases. The Beijing Clade A is associated with XPTB, while 4.3 sublineage is associated with PTB cases (FET Beijing A vs. 4.3: p = 0.0005944). A similar positive association was observed for XPTB with *M.tuberculosis* sublineages Beijing CAO and 4.3, while subclade 4.3 was strongly associated with PTB (Beijing CAO vs. 4.3: p=0.01109).

**Table 1.** Genetic clades, identified among *M. tuberculosis* isolates from different localizations (see fig. 2)

| Genotype | PTB |  | XPTB |  | Total |  |
| --- | --- | --- | --- | --- | --- | --- |
|  | N | % | N | % | N | % |
| Beijing: | 48 | 66.67 | 60 | 82.19 | 108 | 74.48 |
| Beijing unclustered | 16 | 22.22 | 15 | 20.55 | 31 | 21.38 |
| Beijing B0/W148 | 26 | 36.11 | 23 | 31.51 | 49 | 33.79 |
| Beijing Clade A | 3 | 4.17 | 14 | 19.18 | 17 | 11.72 |
| Beijing CAO | 3 | 4.17 | 8 | 10.96 | 11 | 7.59 |
| 4.1 | 5 | 6.94 | 3 | 4.11 | 8 | 5.52 |
| 4.2 | 6 | 8.33 | 4 | 5.48 | 10 | 6.9 |
| 4.3 | 11 | 15.27 | 2 | 2.74 | 13 | 8.97 |
| 4.4 | 1 | 1.39 | 0 | 0 | 1 | 0.69 |
| 4.8 | 1 | 1.39 | 4 | 5.48 | 5 | 3.45 |

Phylogenetic analysis (fig. 2) allowed to discriminate *M.tuberculosis* genetic clusters and to screen for synapomorphic SNVs, Indels that were specific for more than 95% of samples from a particular cluster. Using this approach to statistical analysis did not identify specific SNVs, InDels signatures associated with TB tissue localization across different *M.tuberculosis* phylogenetic groups (Supplementary figure1).

*M.tuberculosis* drug susceptibility testing allowed the recognition of five clinical groups of isolates: (a) susceptible; (b) mono-resistant, i.e., resistant to one drug; (c) poly-resistant – resistant to more than one drug, but not MDR; (d) multidrug-resistant (MDR) and (e) extensively drug-resistant (XDR), according to WHO definition (13,14) (Table 2). Among 73 extrapulmonary *M.tuberculosis* isolates 12 (16.44%) were susceptible to all drugs; 3 (4.11%) were mono-resistant; 9 (12.33%) were poly-resistant; 44 (60.27%) were MDR and 5 (6.85%) were XDR. Seven out of 72 pulmonary isolates (9.72%) were susceptible to all drugs; 2 (2.7%) were mono-resistant; 9 – resistant to several drugs, but not MDR; 30 isolates (41.67%) – MDR and 24 (33.33%) were XDR. The prevalence of XDR phenotype was almost 5-fold higher in PTB than was observed in XPTB cases (33% XDR in PTB vs. 6.8% XDR in XPTB; p= 6.133*10^-05^; Table 2). The majority of bacterial isolates in Beijing cluster were resistant to at least one drug.

**Table 2.**
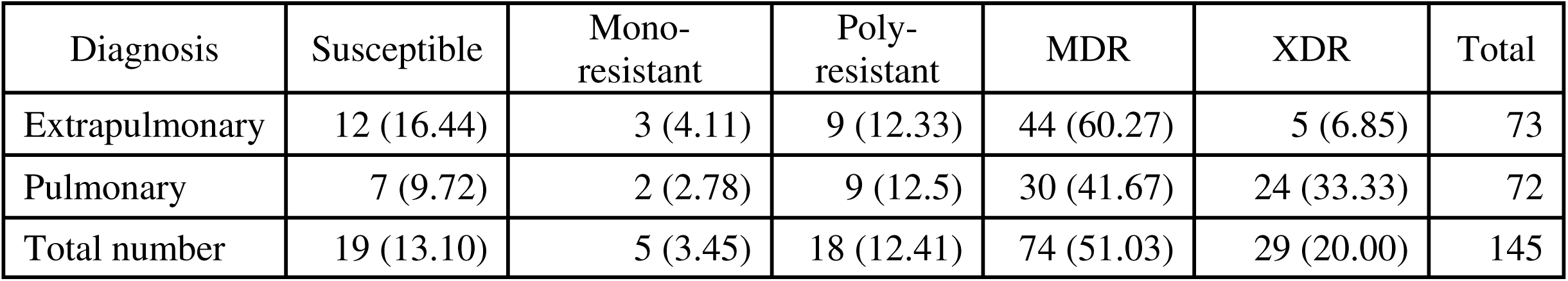
Characterization of drug susceptibility of *M. tuberculosis* from different localizations, n (%)

Over 80% of *M.tuberculosis* isolates in Beijing cluster had MDR (n=62) and XDR (n=25). Only 9 out of 108 isolates were susceptible to all drugs (Table 3). Statistical analysis showed an association of Beijing cluster with drug resistance (p= 0.008542) and especially with M/XDR (p= 3.007*10^-5^) compared to non-Beijing clades.

**Table 3.**
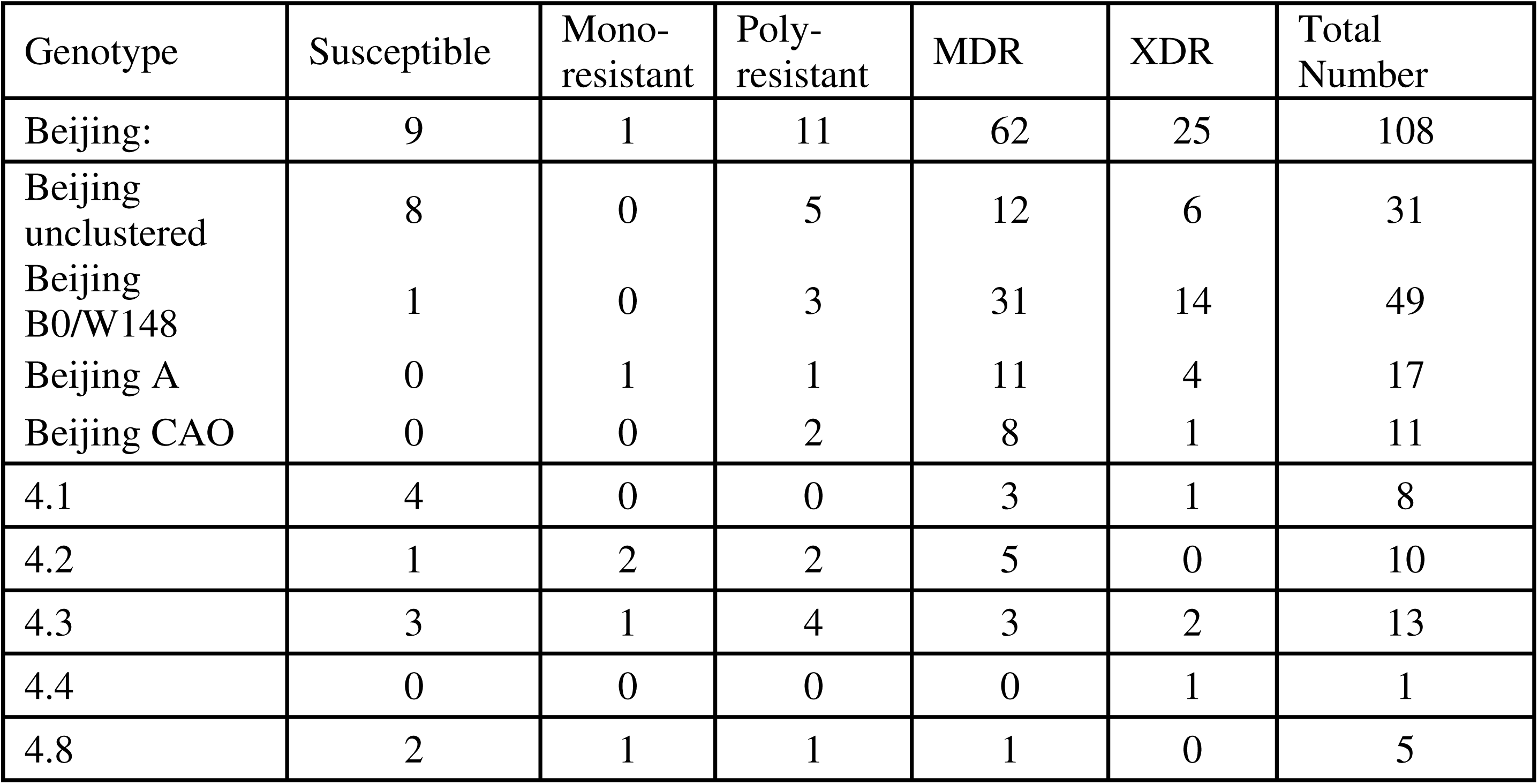
Characterization of drug susceptibility in different *M. tuberculosis* genetic clades (n).

### Mutations associated with drug resistance

WGS data were screened for previously published mutations associated with resistance to TB drugs (15): SM, INH, RIF, OFL, PZA, EMB, ETH, KM, AM and CS (ST2, ST3).

INH-resistance has been associated with mutation S315T in *katG* gene, and 95.87 % of INH-resistant strains in this study carried mutations in these regions. RIF-resistance was associated with mutations in *rpoB* 81-bp core region, 96.15% of resistant isolates carried mutations in this genome region. Over 98% of SM-resistant isolates had mutations in *rpsL, rrs* and *gid* genes. Mutations in promoter region of *eis* gene were identified in 18 KM-resistant and 13 KM-susceptible isolates. It was previously shown that mutations in regulatory region of *whiB7* gene can indirectly influence KM-resistance, however more likely are associated with SM-resistance (16). We identified only two KM-resistant and 3 susceptible isolates with mutations in the region upstream *whiB7* and all of these mutants were resistant to SM. The majority of OFL-resistant isolates (90.48%) carried mutations in *gyrA* or *gyrB* quinolone resistance-determining regions. Mutations in *embB* gene between codones 296 and 497 (17) were detected in 63 resistant and 27 susceptible to EMB isolates. Mutations in *embC*-*embA* intergenic region (8-16 nucleotides upstream *embA* gene) were detected in 16 EMB-resistant and 7 EMB-susceptible isolates. Point mutations were detected in *pncA* and *rpsA* genes in 31 PZA-resistant isolates (67.39%).

A list of SNVs that might be associated with PZA-resistance in *pncA* gene published by Yadon A. et al., 2017 (18) were subjects for discovery in *M.tuberculosis* WGS data. We identified 41 mutations leading to amino acid substitutions in 62 *M.tuberculosis* isolates. Twenty-nine PZA-resistant isolates (63.04%) carried mutations in *pncA* gene, however three of these mutations were previously identified as associated with PZA-susceptibility and six mutations were not related to susceptibility or resistance. Nine PZA-susceptible isolates had SNVs in *pncA* gene; six of these mutations were associated with *in vivo* or/and *in vitro* resistance to PZA and three did not have association with PZA-susceptibility or resistance, according to Yadon et al. (18). Among *M.tuberculosis* isolates with unknown PZA-resistance status (n=24), 14 mutations associated with PZA resistance in 16 *M.tuberculosis* isolates were detected, one mutation in associated with PZA-susceptibility and 6 mutations in 8 genomes with unknown association with resistance or susceptibility to PZA (ST3).

ETH-resistant isolates are known to have mutations located in *ethA/R* locus, *inhA* gene and its promoter (15). Forty-eight sequenced isolates were known to be resistant to ETH. In our dataset, ETH-resistant isolates did not have mutations in *ethA* gene; mutations in *fabG–inhA* operon (−15 and −34 nt) were detected in 9 ETH-resistant and 5 ETH-susceptible isolates. Mutations in *inhA* gene were relatively rare and were detected in 2 resistant and 4 susceptible isolates. Mutations in *ethA* gene were identified in 8 ETH-resistant, 25 ETH-susceptible and 3 isolates with unknown ETH susceptibility data. Mutation in *ethR* gene (A70T) was detected only in one ETH-resistant isolate, while mutations in *ethA-ethR* intergenic region (7 bp upstream *ethA* or 69 bp upstream *ethR* start-codons) was detected in 9 isolates and only three of them were ETH-resistant.

We compared mutations, associated with *M.tuberculosis* drug resistance, detected in our study with a list of mutations that are widely used in molecular-genetic tests for *M.tuberculosis* identification and drug resistance prediction: HAIN GenoType MTBDRplus and GenoType MTBDRsl v 1.0 and v 2.0. The majority of mutations associated with INH-resistance that were detected in our study could be detected by HAIN assay, however mutation in −34 position upstream *fabG-inhA* operon is not included in HAIN catalogue may be due to relatively rare occurrence (2 isolates in our dataset). Detection of mutations associated with RIF-resistance using HAIN assay would also allow to identify most of RIF-resistant isolates in our dataset, however Q432K mutation in *rpoB* gene detected in one RIF-resistant strain and might be related with RIF-resistance. Mutations associated with EMB-resistance detected by HAIN diagnostic assay are represented by M306V and M306I substitutions only. In our study M306V was the most frequent mutation (n=20), M306I substitution were identified in 5 genomes, while 38 *M.tuberculosis* isolates that had mutations in *embB* region between codons 296 and 497 that could not be detected by HAIN system. Mutations in *embA* promoter region that are not represented in HAIN test-system were detected in 14 EMB-resistant isolates. It should be noted that mutations in *embB* gene and *embA* promoter associated with EMB-resistance often detected among EMB-susceptible isolates. For instance, mutation M306V in *embB* gene was detected in 20 EMB-resistant and 9 EMB-susceptible isolates. Mutations associated with resistance to aminoglycosides and peptide antibiotics are represented in HAIN test by 3 mutations in *rrs* gene, associated with KM/AM/CM/viomycin resistance, and *eis* promoter mutations, associated with KM resistance. We did not find mutations in *rrs* positions 1401, 1402 and 1484, but identified mutation in 1490 position in one CM-resistant isolate. Three out of four SNVs identified in *eis* promoter could be detected by HAIN assay (10, 12 and 14 bp upstream *eis* start codon) however in our study these mutations were detected in 18 KM-resistant and 12 KM-susceptible isolates. GenoType MTBDRsl assay allows to detect mutations in *gyrA* 90, 91 and 94 codons and in *gyrB* 538 and 540 codons, associated with OFL-resistance. In our dataset most of OFL-resistance mutations were detected in *gyrA* gene and correspond to the variety of mutations presented in the HAIN system, though mutations in *gyrB* gene that were detected in several OFL-resistant strains were not covered by HAIN line-probe assay.

### TB/HIV co-infection

Bacterial isolates were collected from 120 HIV-negative patients and 25 HIV-infected patients. We observed a higher rate of generalized TB among HIV-infected individuals (Table 4) (p= 6.579*10^-05^). XPTB was significantly more frequent among HIV-positive patients compared to patients with PTB (Table 4; p=0.000998). Thus, HIV-infection increases the probability of TB generalization or development of active XPTB. XPTB development was significantly higher among patients infected with HIV carrying the Beijing-unclustered strain of *M.tuberculosis* (p=0.068) (Table 5). A similar XPTB excess was apparent among patients carrying the Beijing B0/W148 strain (p= 0.0013) (Table 5). Other genetic groups with HIV co-infection did not make a significant impact on pulmonary or extrapulmonary TB development (Table 5).

**Table 4.** Number of TB/HIV co-infection cases.

| Diagnosis | HIV+ | HIV- | N |
| --- | --- | --- | --- |
| PTB | 3 | 69 | 72 |
| XPTB | 14 | 41 | 55 |
| Generalized TB | 8 | 10 | 18 |

**Table 5.** Number of *M. tuberculosis* isolates obtained from patients with different TB localization and HIV-status.

| Geneine clage | HIV co-infection | Diagnosis |  |
| --- | --- | --- | --- |
|  |  | PTB | XPTB |
| 4.8 | + | 0 | 1 |
|  | - | 1 | 3 |
| 4.3 | + | 1 | 1 |
|  | - | 10 | 1 |
| 4.2 | + | 1 | 0 |
|  | - | 5 | 4 |
| 4.1 | + | 0 | 0 |
|  | - | 5 | 3 |
| 4.4 | + | 0 | 0 |
|  | - | 1 | 0 |
| Beijing unclustered | + | 0 | 6 |
|  | - | 16 | 9 |
| Beijing B0/W148 | + | 1 | 10 |
|  | - | 25 | 13 |
| Beijing CAO | + | 0 | 1 |
|  | - | 4 | 7 |
| Beijing A | + | 0 | 3 |
|  | - | 2 | 11 |
| Total |  | 72 | 73 |

## Discussion

Our study is devoted to the comparative analysis of *M.tuberculosis* isolates obtained from patients with different clinical features of the disease – pulmonary and extrapulmonary TB. Our previous analysis of *M.tuberculosis* isolates from patients with tuberculous spondylitis showed that this group of bacterial strains is characterized by low genetic diversity and high prevalence of Beijing isolates (82% of strains belonged to Beijing clade) (3). Prevalence of Beijing genetic group among extrapulmonary *M.tuberculosis* isolates compared to pulmonary ones was also reported in a recent study, which identified Beijing in 75% of the extrapulmonary *M.tuberculosis* cases (19). Our current study showed that extrapulmonary *M.tuberculosis* isolates belong to Beijing genetic group (82.2%) more often than pulmonary (66.7%). Despite there was a higher prevalence of Beijing strains among extrapulmonary *M.tuberculosis* samples (82.19% vs 66.67%), in our study pulmonary strains were frequently associated with XDR (p= 6.133*10^-05^). It was previously shown that *M.tuberculosis* strains from Beijing genetic group are highly associated with M/XDR (20-22). Our results confirm this statement: the frequency of drug resistance among *M.tuberculosis* Beijing isolates was higher compared to other genetic groups in both pulmonary and extrapulmonary strains (Table 1). *M.tuberculosis* genetic lineage 4 identified in our study was also previously detected in Russia: Ural family (4.2) identified in many regions in Eurasia (23), and heterogeneous sublineage 4.1 and sublineage 4.3 were observed in many countries across the world (23). Like 4.2 sublineage 4.4 occurred in high proportions among isolates from particular countries in Asia and Africa, but were largely absent from the Americas. Sublineage 4.8 was identified as a part of the 4.10 genetic group, the latter was previously mentioned as one of the most widely spread sub-clades of the lineage 4. (24).

It was shown that Beijing A and CAO isolates are significantly more often identified in extrapulmonary TB cases, while isolates from principal genetic lineage 4.3 were mostly identified in patients with pulmonary TB (Table 1). Association of specific genetic clades and sub-clades of *M.tuberculosis* strains with TB diagnosis, shown on our dataset, is an interesting fact which suggests some specific molecular signatures in different *M.tuberculosis* genetic groups that can make an impact on active TB disease localization at least in Russian population of TB patients.

Analysis of mutations, associated with susceptibility to first and second line TB drugs arevealed that relatively high proportion of INH-, RIF-, SM- and OFL- resistant isolates had standard SNVs predictive of drug-resistance. However, some SNVs, associated with resistance to TB drugs cannot be unequivocally interpreted. For example, the proportion of EMB-resistant isolates with mutations in *embB* gene between codones 296 and 497 and *embC-embA* intergenic region were identified in 94.29% of EMB-resistant isolates, however over 40% of EMB susceptible isolates carried the same mutations. Similar situation was revealed with mutations, associated with KM-resistance in *eis* and *whiB7* promoter regions – 42.5% of KM-resistant and 22.4% of KM-susceptible isolates carried mutations in these regions (ST3). This observation might be related with the fact that mutations can be responsible for the development of high and low levels of drug resistance, while in our study we did not have data on minimum inhibitory concentration. *M.tuberculosis* isolates from our dataset were checked for mutations in *pncA* and *rpsA* genes that were known to be involved in development of resistance to PZA (17, 25). Only 4 PZA-resistant isolates carried mutations in *rpsA* gene, but the majority of PZA-resistant isolates had mutations in *pncA* gene mentioned in previously published catalog of mutations associated with susceptibility to PZA (17). Despite most of obtained data were consistent with previously published data, in PZA-resistant strains we detected few mutations, that were mentioned as associated with susceptibility to PZA and there were several PZA-susceptible isolates, that carried mutations associated with PZA-resistance according to Yadon et al. data. This fact may indicate either that these mutations are not involved into the development of PZA resistance, or their influence on resistance to higher doses of the drug that were analyzed in our research. Detection of mutations associated with drug resistance among susceptible isolates may indicate the presence of low number of drug resistant clones in *M.tuberculosis* population and might be a signal for correction of TB therapy in case it was developed based on phenotypic data.

Comparative analysis of *M.tuberculosis* isolates obtained from HIV-positive and HIV-negative patients would suggest that HIV co-infection increases the probability of TB generalization or development of XPTB. Previous epidemiological studies showed similar results – a number of studies have identified a relationship between HIV and the development of XPTB (26-28). Results of the recent research showed that HIV infection is one of risk factors for XPTB in the US (29). The majority of *M.tuberculosis* strains included in our study (both pulmonary and extrapulmonary) belonged to Beijing genetic group widely spread in the Russia. Comparative analysis of XPTB and PTB cases in HIV-positive and HIV-negative patients infected with *M.tuberculosis* strains from various genetic groups allowed to obtain produced different results. For example, the HIV correlates with XPTB in patients infected with Beijing B0/W148 and unclustered Beijing strains. However, HIV co-infection is not associated with XPTB in patients infected with bacterial strains related to genetic clades Beijing CAO, A and 4.8 (associated with XPTB) and strains from 4.3 clade (associated with PTB). Thus, according to our data, it may be possible to assess the risk of XPTB or generalized TB, depending on the pathogen strain and the presence of HIV co-infection. A limitation of our study was a relatively low number of *M.tuberculosis* isolates obtained from HIV-infected individuals. We suggest that further analysis of TB/HIV co-infection with attention on *M.tuberculosis* phylogeny could help better understanding processes leading to active XPTB and generalized TB development.

XPTB development is a multifactorial process that can depend on many factors, including the biology of the pathogen and the host. Understanding of these factors can help in assessing the risks of developing complicated forms of TB, optimizing diagnosis and treatment. Our research allowed making a snapshot of genomic markers identified in *M.tuberculosis* strains obtained from patients with PTB and XPTB in Russia. Further comprehensive analysis of bacterial and human biological signatures might allow for better understanding consistent pattern of XPTB development.

## Acknowledgements

Scientific research was performed using equipment of the Resource Center «Biobank», Research park of St. Petersburg State University.

## References

1. GLOBAL TUBERCULOSIS REPORT 2017. World Health Organization; 2017 p. 187.

2. Yablonsky P, Mushkin Ay, Belilovsky E, Galkin V. Extrapulmonary tuberculosis. In: Tuberculosis in the Russian Federation, 2012/2013/2014 Analytical review of statistical indicators for tuberculosis used in the Russian Federation and in the world. p. 129–35.

3. Chernyaeva E, Rotkevich M, Krasheninnikova K, et al. Whole-Genome Analysis of Mycobacterium tuberculosis from Patients with Tuberculous Spondylitis, Russia. Emerging Infectious Diseases. 2018;24(3):-583. doi:10.3201/eid2403.170151.

4. World Health Organization. Guidelines for surveillance of drug resistance in tuberculosis. Geneva: WHO; 2009.

5. van Embden JD, Cave MD, Crawford JT, Dale JW, Eisenach KD, Gicquel B, et al. Strain identification of Mycobacterium tuberculosis by DNA fingerprinting: recommendations for a standardized methodology. J Clin Microbiol. 1993 Feb;31(2):406–9.

6. Bolger AM, Lohse M, Usadel B. Trimmomatic: a flexible trimmer for Illumina sequence data. Bioinforma Oxf Engl. 2014 Aug 1;30(15):2114–20.

7. Stamatakis A. RAxML-VI-HPC: maximum likelihood-based phylogenetic analyses with thousands of taxa and mixed models. Bioinformatics. 2006;22:2688–90

8. Benavente ED, Coll F, Furnham N, McNerney R, Glynn JR, Campino S, et al. PhyTB: Phylogenetic tree visualisation and sample positioning for M. tuberculosis. BMC Bioinformatics [Internet]. 2015 Dec [cited 2016 Nov 10];16(1).

9. Xia E, Teo YY, Ong RT. SpoTyping: fast and accurate in silico Mycobacterium spoligotyping from sequence reads. Genome Med. 2016 Feb 17;8(1):19. doi: 10.1186/s13073-016-0270-7. PubMed PMID: 26883915; PubMed Central PMCID: PMC4756441.

10. Yin Q, Liu H, Jiao W, Li Q, Han R, Tian J, et al. Evolutionary History and Ongoing Transmission of Phylogenetic Sublineages of Mycobacterium tuberculosis Beijing Genotype in China. Sci Rep. 2016 Sep 29;6:34353.

11. Mokrousov I. Insights into the Origin, Emergence, and Current Spread of a Successful Russian Clone of Mycobacterium tuberculosis. Clin Microbiol Rev. 2013 Apr 1;26(2):342–60.

12. Shitikov E, Kolchenko S, Mokrousov I, Bespyatykh J, Ischenko D, Ilina E, et al. Evolutionary pathway analysis and unified classification of East Asian lineage of Mycobacterium tuberculosis. Sci Rep [Internet]. 2017 Dec [cited 2017 Oct 30];7(1). Available from: http://www.nature.com/articles/s41598-017-10018-5

13. What is multidrug-resistant tuberculosis (MDR-TB) and how do we control it? [Internet]. 2017. Available from: http://www.who.int/features/qa/79/en/

14. Drug-resistant TB: XDR-TB FAQ. Available from: http://www.who.int/tb/areas-of-work/drug-resistant-tb/xdr-tb-faq/en/

15. Zhang Y, Yew W-W. Mechanisms of drug resistance in Mycobacterium tuberculosis: update 2015. Int J Tuberc Lung Dis Off J Int Union Tuberc Lung Dis. 2015 Nov;19(11):1276–89.

16. Reeves AZ, Campbell PJ, Sultana R, Malik S, Murray M, Plikaytis BB, et al. Aminoglycoside cross-resistance in Mycobacterium tuberculosis due to mutations in the 5’ untranslated region of whiB7. Antimicrob Agents Chemother. 2013 Apr;57(4):1857–65.

17. Yakrus MA, Driscoll J, McAlister A, Sikes D, Hartline D, Metchock B, et al. Molecular and Growth-Based Drug Susceptibility Testing of *Mycobacterium tuberculosis* Complex for Ethambutol Resistance in the United States. Tuberc Res Treat. 2016;2016:1–5.

18. Yadon AN, Maharaj K, Adamson JH, Lai Y-P, Sacchettini JC, Ioerger TR, et al. A comprehensive characterization of PncA polymorphisms that confer resistance to pyrazinamide. Nat Commun. 2017 198(1):588.

19. Vyazovaya A, Mokrousov I, Solovieva N, Mushkin A, Manicheva O, Vishnevsky B, et al. Tuberculous Spondylitis in Russia and Prominent Role of Multidrug-Resistant Clone Mycobacterium tuberculosis Beijing B0/W148. Antimicrob Agents Chemother. 2015 Apr;59(4):2349–57.

20. Couvin D, Rastogi N. Tuberculosis - A global emergency: Tools and methods to monitor, understand, and control the epidemic with specific example of the Beijing lineage. Tuberc Edinb Scotl. 2015 Jun;95 Suppl 1:S177–189.

21. Chernyaeva E, Fedorova E, Zhemkova G, Korneev Y, Kozlov A. Characterization of multiple and extensively drug resistant Mycobacterium tuberculosis isolates with different ofloxacin-resistance levels. Tuberculosis. 2013 May;93(3):291–5.

22. de Steenwinkel JEM, ten Kate MT, de Knegt GJ, Kremer K, Aarnoutse RE, Boeree MJ, et al. Drug Susceptibility of *Mycobacterium tuberculosis* Beijing Genotype and Association with MDR TB. Emerg Infect Dis. 2012 Apr;18(4):660–3.

23. Mokrousov I. Mycobacterium tuberculosis phylogeography in the context of human migration and pathogen’s pathobiology: Insights from Beijing and Ural families. Tuberculosis. 2015 Jun;95:S167–76.

24. Stucki D, Brites D, Jeljeli L, Coscolla M, Liu Q, Trauner A, et al. Mycobacterium tuberculosis lineage 4 comprises globally distributed and geographically restricted sublineages. Nat Genet. 2016 Dec;48(12):1535–43.

25. Liu W, Chen J, Shen Y, Jin J, Wu J, Sun F, Wu Y, Xie L, Zhang Y, Zhang W. Phenotypic and genotypic characterization of pyrazinamide resistance among multidrug-resistant Mycobacterium tuberculosis clinical isolates in Hangzhou, China. Clin Microbiol Infect. 2017 Dec 26. pii: S1198-743X(17)30687-0. doi: 10.1016/j.cmi.2017.12.012.

26. Desforges JF, Barnes PF, Bloch AB, Davidson PT, Snider DE. Tuberculosis in Patients with Human Immunodeficiency Virus Infection. N Engl J Med. 1991 Jun 6;324(23):1644–50.

27. Slutsker L, Castro KG, Ward JW, Dooley SW. Epidemiology of Extrapulmonary Tuberculosis Among Persons with AIDS in the United States. Clin Infect Dis. 1993 Apr 1;16(4):513–8.

28. Naing C, Mak JW, Maung M, Wong SF, Kassim AIBM. Meta-Analysis: The Association Between HIV Infection and Extrapulmonary Tuberculosis. Lung. 2013 Feb;191(1):27–34.

29. Qian X, Nguyen DT, Lyu J, Albers AE, Bi X, Graviss EA. Risk factors for extrapulmonary dissemination of tuberculosis and associated mortality during treatment for extrapulmonary tuberculosis. Emerg Microbes Infect. 2018 Jun 6;7(1):102.

